# Uncovering the molecular signatures of russet skin formation in Niagara grapevine (*Vitis vinifera x Vitis labrusca*)

**DOI:** 10.1101/2023.08.25.554330

**Authors:** Guilherme Francio Niederauer, Geovani Luciano de Oliveira, Alexandre Hild Aono, Diego da Silva Graciano, Sandra Maria Carmello-Guerreiro, Mara Fernandes Moura, Anete Pereira de Souza

## Abstract

Grape breeding programs are mostly focused on developing new varieties with high production volume, sugar contents, and phenolic compound diversity combined with resistance and tolerance to the main pathogens under culture and adverse environmental conditions. The ‘Niagara’ variety (*Vitis labrusca* x *Vitis vinifera*) is one of the most widely produced and commercialized table grapes in Brazil. In this work, we selected three Niagara somatic variants with contrasting berry phenotypes and performed morphological and transcriptomic analyses of their berries. Histological sections of the berries were also performed to understand anatomical and chemical composition differences of the berry skin between the genotypes. An RNA-Seq pipeline was implemented, followed by global coexpression network modeling. ‘Niagara Steck’, an intensified russet mutant with the most extreme phenotype, showed the largest difference in expression and showed selection of coexpressed network modules involved in the development of its russet-like characteristics. Enrichment analysis of differently expressed genes and hub network modules revealed differences in transcription regulation, auxin signaling and cell wall and plasmatic membrane biogenesis. Cutin- and suberin-related genes were also differently expressed, supporting the anatomical differences observed with microscopy.

## Introduction

Grapevine (*Vitis spp*.) is one of the most important fruit crops worldwide in terms of hectares cultivated and economic value. In Brazil, the American *Vitis labrusca* varieties and hybrids (*V. labrusca* x *V. vinifera*) are successful due to their improved adaptation to the high humidity of the country. These varieties predominate in Brazilian plantations, presenting voluminous production due to their greater hardiness, such as tolerance to the main diseases that attack grapevine under these conditions^1^. Among these hybrids, the Niagara variety stands out as one of the main table grapes cultivated and sold in Brazil since its addition to the country’s vine crops in the 19th century.

The ‘Niagara Rosada’ variety (N. Rosada, *V. labrusca* x *V. vinifera*) is the most predominant of the group, and it is traditionally cultivated in the southeastern and southern regions of the country. This variety arose due to a natural somatic mutation that occurred in the ‘Niagara Branca’ variety (N. Branca), differing mainly in terms of the color of the berries. The skin of N. Rosada berries has a strong pink tone, while in N. Branca, a light green tone is observed. These two varieties have characteristics that are appreciated by the Brazilian market, such as mucilaginous pulp, a raspberry flavor, low acidity and full medium-sized bunches^2^. In addition to N. Branca and N. Rosada, other mutants of the Niagara genotype were identified, the most distinct of which is ‘Niagara Steck’ (N. Steck). This variety displays irregular cluster development resulting in unusual, intensified russet characteristics in grapes, as evidenced by the berry skin, which is rough, thick and rust-colored. The pulps of the berries are also affected, being smaller and more acidic than those of the other Niagara varieties.

A soft texture and specific pigmentation of grape berry skin are among the attributes most valued by consumers and are driven by the breakdown of skin cell walls and anthocyanin accumulation in colored varieties, respectively. Wax production and the cuticular layer also contribute to berry skin attractiveness by conferring a lustrous and glossy appearance. In this sense, disruptions and alterations in the grape berry skin of a cultivar have a great impact on its market acceptance and are treated with great attention. The berry characteristics of N. Steck are observed independently of environmental conditions and agronomic practices and have been observed yearly since its inclusion in the germplasm bank^3^. Despite the evident phenotypic differences, identical molecular profiles were observed between these three Niagara varieties using highly polymorphic microsatellite markers^4^.

Grape berry quality is assessed by numerous types of measurements, such as phenolic composition, skin color, texture and thickness, berry size, chunk size and weight, berry count per chunk, Brix degree, pulp volume and acidity and aromatic composition of fresh and processed berries. The development of these attributes starts after the initial structural formation of the berry, 0 to 40 days after flowering (DAF), and it is marked by the initiation of skin pigmentation and sugar accumulation (veraison, 60 DAF), followed by berry swelling and skin softening, until the full maturation (ripe) stage is reached (90-120 DAF)^5^. These stages have been widely studied, and many of the main hormones, genes, and transcription factors involved in their regulation have already been identified^6–8^. Recent grapevine transcriptomic studies have provided a better understanding of the regulatory switches controlling the phenylpropanoid pathway and how they are influenced by somatic mutations and environmental and biotic stressors by means of hormonal feedbacks, coexpression networks and epigenetic modifications, resulting in an extensive variety of phenotypes and resistance mechanisms in grape cultivars^9–11^. In this context, global analysis of the expression of millions of transcripts simultaneously, through RNA-Seq accompanied by coexpression network analysis, has the potential not only to identify gene expression patterns with a high level of accuracy and at great depth but also to reveal correlations between these patterns and how they regulate the stages of berry development, thus allowing a better understanding of the role of specific genes and gene interactions in determining phenotypic diversity^12^.

The berries of ‘Niagara’ grape mutants differ in several characteristics, many of which have economic potential for Brazilian viticulture. This set of changes shows that phenotypic variations in grapes are not derived from a single gene but from interactions of genes in a network. To understand how these characteristics are manifested, identifying the biological pathways involved is of great importance, enabling the prospection of economically important genes within grapevine breeding programs. Although ‘Niagara’ is one of the main grape cultivars in Brazil, to date, no studies have investigated the expression patterns related to the development of its berries. Accounting for the genomic similarities between the mentioned varieties, the transcriptomic and coexpression approaches, applied in combination with contrasting phenotypes such as those in N. Steck, can aid in the identification of novel and specific genes that influence the phenotypic determination of grapevine berries.

In the current study, we utilized berry-focused transcriptomic analysis coupled with coexpression network modeling to investigate genes and pathways controlling the maturation of berries in three contrasting varieties of the Niagara grapevine cultivar. We performed anatomical analysis of the skin, differential expression analysis by means of RNA-Seq at two distinct stages of berry development (*veraison* and complete maturation), and coexpression network modeling through a scale-free topology approach.

## Results

### Berry Skin Structural Analysis

#### Structural Variations

Since the sampled varieties exhibit different phenotypic profiles, we asked whether such alterations are evidenced in the structure of the skin in addition to the contrasting pigmentation. When comparing the anatomical structure of the skin region between the three studied genotypes, we observed that in the varieties N. Branca and N. Rosada, at both development stages, the epicarp comprises a uniseriate epidermis covered by cuticle, with small, rectangular, compactly arranged cells consisting of thick cell walls (Fig. 1a-f). Furthermore, in the second maturation phase of these two varieties, different patterns of epicuticular wax deposition occurred (Fig. 1c, f).

**Figure 1.**
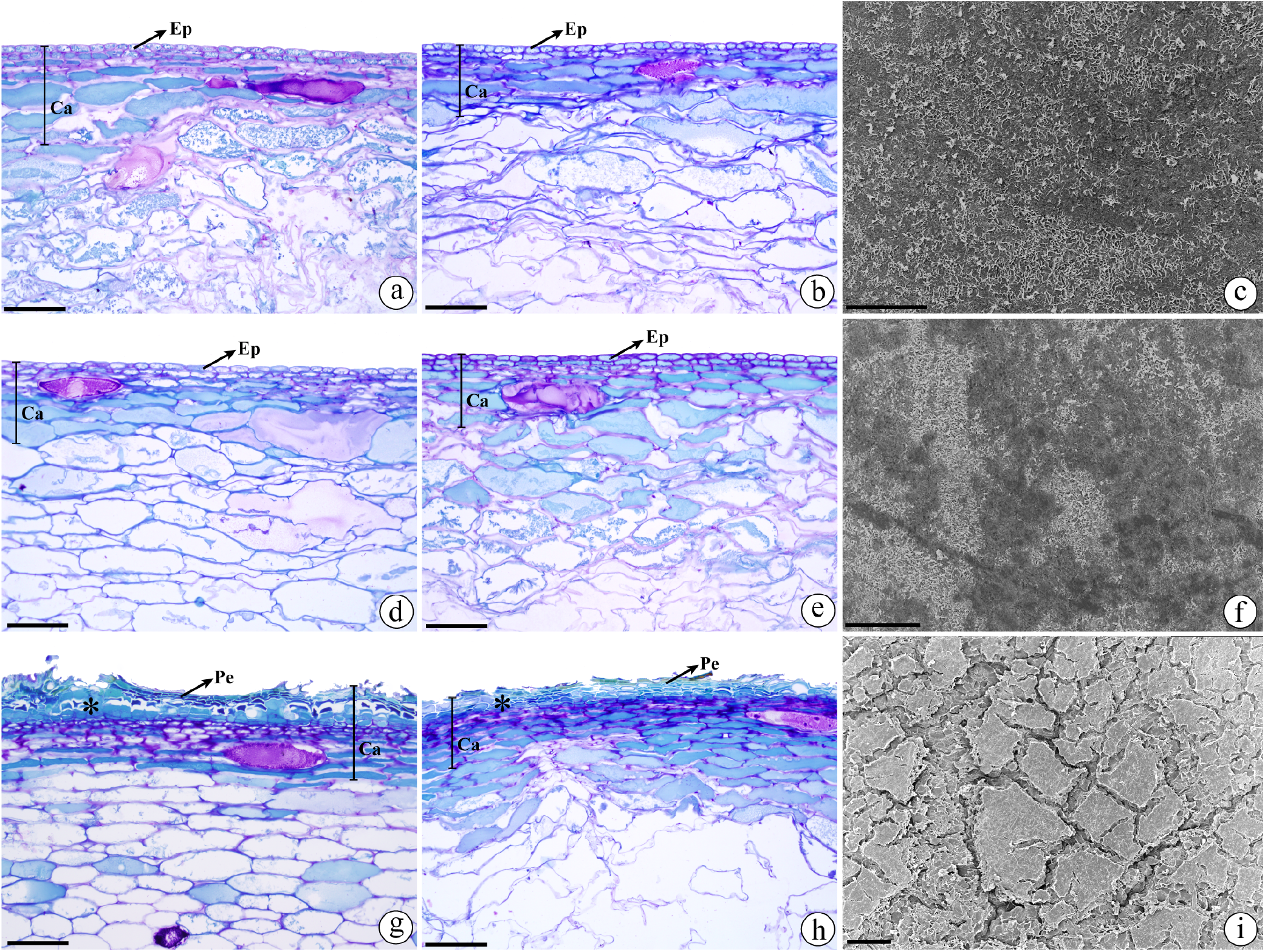
Anatomical structure and micromorphology of the skin of different Niagara varieties. a) N. Branca - Veraison. (b) N. Branca - Maturation. (c) SEM surface of the epicarp of N. Branca - Veraison. (d) N. Rosada - Veraison. (e) N. Rosada - Maturation. (f) SEM surface of the epicarp of N. Rosada - Maturation. (g) N. Steck - Veraison. (h) N. Steck - Maturation. (i) SEM surface of the epicarp of N. Steck Genotype - Maturation. Asterisk indicates the region of lateral meristem that forms the periderm. Abbreviations: Ca, skin; Ep, epidermis; Pe, periderm. Scale bars: 50 *μ*m (c, f); 100 *μ*m (a-b, d-e, g-i).

Distinctively, the epicarp of the N. Steck genotype consists of a periderm that covers the surface of the pericarp with several cell layers (Fig. 1g-h). In this case, the epidermis is collapsed, giving way to scar tissue that develops by periclinal divisions of cells from the subepidermal layers of mesocarp origin (Fig. 1g, i). A portion of the outer mesocarp together with the epicarp composes the skin in the three varieties. In this region, the mesocarp is of parenchymal origin, presenting two or three layers of small and rectangular cells under the epicarp constituting the subepidermal layers (Fig. 1a-b, d-e, g-h). Underlying these layers, the outer mesocarp consists of large, tangentially elongated cells with phenolic or, less frequently, mucilaginous content, in addition to large intercellular spaces (Fig. 1a-b, d-e, g-h). The cells of the inner mesocarp disintegrate as a result of the fruit’s maturation processes, except in the N. Steck genotype at *veraison*, when the cell walls remain intact (Fig. 1g).

#### Biochemical Composition

To define the profile of the chemical compounds present in the skin, we compared the phenotypes through histochemical analysis. When conducting these tests, we observed the presence of pectic substances in the conspicuous intercellular spaces of the outer mesocarp in the three varieties, with a more intense reaction in the N. Steck genotype, as well as the presence of cells with mucilaginous content (Fig. 2a, b). The tests also revealed the presence of cells with phenolic content in the epicarp and external mesocarp of the three genotypes, with a more intense reaction in the periderm cells of the N. Steck genotype in both maturation phases (Fig. 2). Tests for lipid substances, such as cutin and suberin, indicated the presence of a thick cuticle in the Branca and Rosada varieties at both maturation stages (Fig. 2e, g). For the Steck genotype, the peridermal cells showed a thickened cell wall consisting of lipid material (Fig. 2f, h-i). In this variety, a thin and irregular cuticle layer also remained (Fig. 2h). The specific test for lignified cell wall identification did not detect lignin. These results demonstrate that although the N. Steck genotype does not have the same epicarp structure as the other varieties, the structure is replaced by cell layers impregnated with lipid substances, making these layers impermeable.

**Figure 2.**
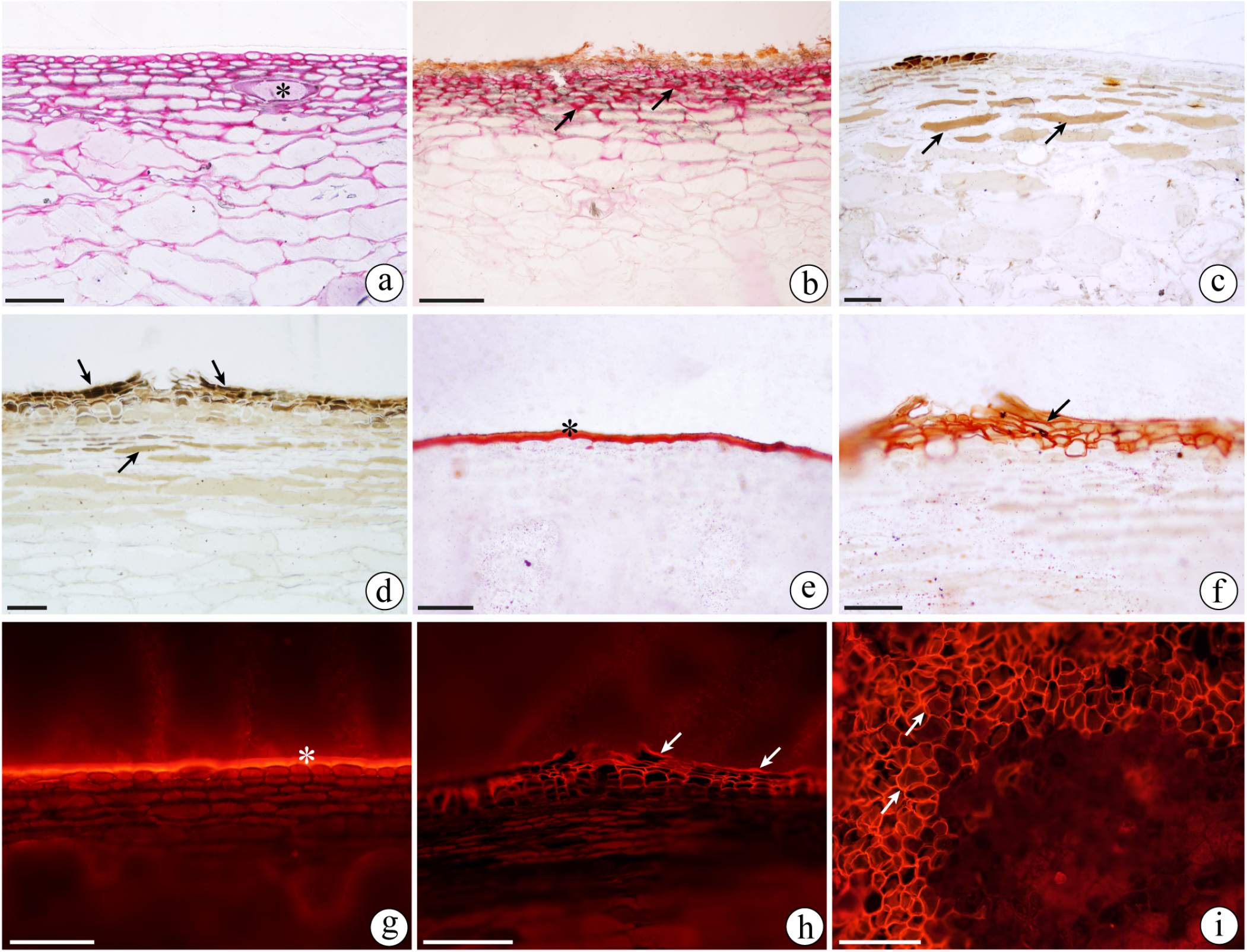
Histochemical tests of cross-sections and paradermal sections of the skin of different Niagara varieties. (a-b) Positive reaction in varieties N. Branca and N. Steck at maturation, respectively, to ruthenium red showing the wide intercellular spaces with pectic substances (arrows) and cells with mucilaginous content (asterisk). (c-d) Positive reaction showing phenolic compounds (arrows) in cells of N. Branca and N. Steck at veraison, respectively. (e) Positive reaction to Sudan III, showing thick cuticle (asterisk) in variety N. Branca at maturation. (f) Positive reaction to Sudan III showing secondary cell walls with lipid substances (arrow) in N. Steck at veraison. (g) Positive reaction to Nile Red fluorochrome showing thick cuticle (asterisk) in the variety N. Rosada at maturation. (h) Positive reaction to Nile Red fluorochrome showing the cell walls with lipid substances of the periderm cells and the cuticle (arrow) in N. Steck at veraison. (i) Paradermal section of the skin region of N. Steck at maturation with a positive reaction to Nile Red fluorochrome, showing the cell walls with lipid substances of the periderm cells. Scale bars: 50 *μ*m (c-e); 100 *μ*m (a-b, f-i).

### Bioinformatics

#### Transcriptome analysis

RNA-Seq libraries generated 423,812,688 reads, of which 5% were removed by the trimming procedure. The mean rate of reads uniquely mapped by STAR against the grapevine reference genome varied between 89.65% and 92.38% (Supplementary Table S1) across all samples. StringTie assembled 76,707 transcripts in total (43,204 unigenes), 75,720 of which were based on reference sequences and 987 novel transcripts (787 unigenes). BUSCO quality assessment revealed the presence of 97.2% of conserved orthologs from the Viridiplantae odb_10 database in the assembled transcriptome (94.1% single and 3.1% duplicated). The read mapping rate against the transcriptome ranged between 87.37% and 89.53% (Supplementary Table S1). Annotations were successfully retrieved for 52.8% of the genes from the Ensemble Plants database and for 32% from sequence alignments with the UniProtKB/Swiss-Prot database (85% of the 43,204 total genes could be annotated). Read count filtering resulted in 19,971 unigenes with expression greater than 10 TMM.

#### Differential Expression Analysis

The differential expression analysis yielded 6,247 unique differentially expressed genes (DEGs) across all contrasts, including comparisons between genotypes and stages within genotypes. For comparisons between genotypes at the veraison stage (Fig. 3A), the NRV vs NBV contrast revealed 919 DEGs (493 more highly expressed in NBV and 426 in NRV), of which only 85 were exclusively observed in this comparison (Fig. 3B). The contrasts including N. Steck at the veraison stage presented a much higher number of DEGs. In contrasts with NBV, we found 2,667 DEGs (967 more highly expressed in NBV and 1,700 more highly expressed in NSV), with 248 exclusive to these contrasts (Fig. 3B). When contrasting NSV and NRV, 3,943 DEGs were found (1,465 more highly expressed in NRV and 2,478 more highly expressed in NSV), with 1,157 of these not reported to be differently expressed in any other comparison of this stage (Fig. 3B).

**Figure 3.**
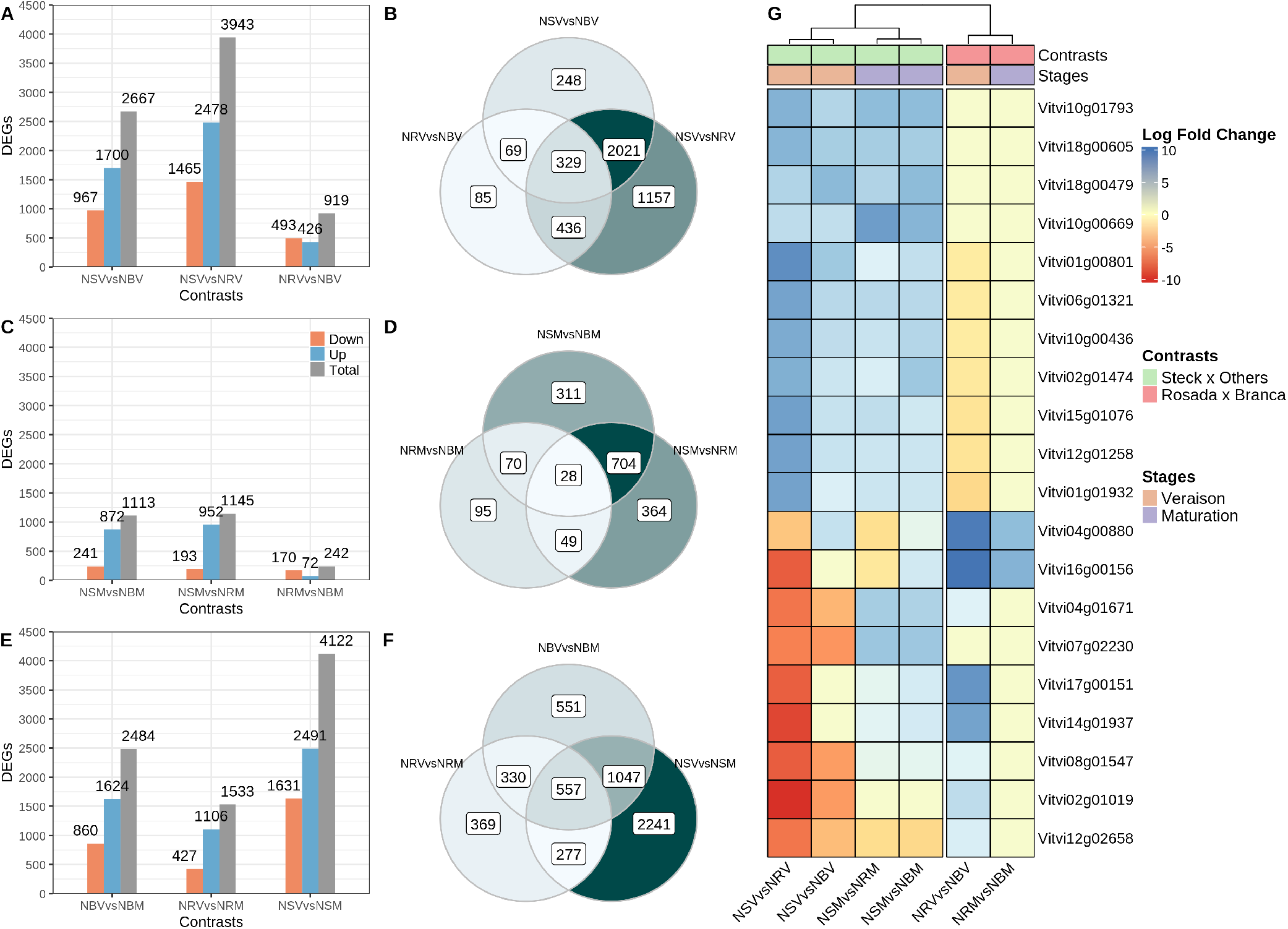
Differentially expressed genes and their correlations with the contrasts. A, B) Veraison. C, D) Maturation. E, F) Same variety across stages. G) Heatmap of the top 20 differentially expressed genes based on log fold change values.

For all comparisons between varieties at the maturation stage (Fig. 3C), a decrease in the number of DEGs was observed, with only 242 between NRM and NBM, of which 95 were only differentially expressed between these two varieties at this stage (Fig. 3D). The N. Steck comparisons yielded fewer than half of the DEGs observed at veraison, with an intersection of 704 DEGs between the NSM vs NBM and NSM vs NRM comparisons.

Considering the comparison of NRV and NBV, there was an apparently balanced proportion of more highly expressed genes (46% and 53%, respectively). Comparisons with NSV revealed a larger proportion of more highly expressed DEGs than in both NBV and NRV. Although the NSV vs NRV contrast showed 1,276 more DEGs than the NSV vs NBV contrast, they showed similar proportions of genes more highly expressed in NSV (62% and 63% for NSV vs NRV and NSV vs NBV, respectively), revealing higher expression of genes in the N. Steck genotype overall. Interestingly, the expression of 1,508 genes was upregulated in NSV compared with both NRV and NBV.

Upon completion of the maturation process, the expression differences between N. Rosada and N. Branca became even smaller; however, there was a larger proportion of more highly expressed DEGs in NBM (70%). The same result was observed in the N. Steck contrasts; there was also a reduction in the total number of DEGs, with an increase in the proportion of DEGs more highly expressed in NSM: 78% of DEGs found in the contrast NSM vs NBM and 83% found in NSM vs NRM.

The expression contrasts of samples from the same variety across stages yielded the most DEGs (Fig. 3E, F). The N. Rosada variety had the fewest DEGs (1,533) in this category, followed by N. Branca (2,484) and N. Steck (4,122), which showed the most DEGs among the nine evaluated expression contrasts. By comparing the DEGs found in the contrasts between stages within varieties, we observed that N. Steck presented the largest number of exclusive DEGs (2,241). Additionally, the intersection of DEGs between N. Steck and N. Branca (1,047 DEGs) was more pronounced than that between N. Steck and N. Rosada (277). For the N. Rosada and N. Branca varieties, we found 369 and 551 exclusive DEGs, respectively, which were notably smaller than the number of DEGs shared between the three varieties across stages (557, Fig. 3F).

Figure 3G highlights the correlation between the 20 top-ranked DEGs according to the fold-change values and the genotype contrasts, revealing very similar patterns of gene expression between N. Rosada and N. Branca, with the exception of the Vitvi04g00880 (GST4) and Vitvi16g00156 (UFGT) genes, which are directly related to the biosynthesis and accumulation of anthocyanins in the grape berry^7,13^, as well as Vitvi17g00151 (malate synthase), a known mediator of fruit and grape ripening^14^, showing upregulated expression in N. Rosada. In the heatmap, all downregulated genes in the N. Steck contrasts showed a decrease in log fold change from veraison to maturation and, in some cases, even became upregulated compared with their expression in both N. Rosada and N. Branca (e.g., Vitvi04g01671 and Vitvi07g02230). The same pattern was not observed for the upregulated genes, which, in most cases, remained upregulated even after the berries reached full ripeness (e.g., Vitvi10g01793, Vitvi18g00805, and Vitvi18g00479). Notably, the VvMYBA1 gene (Vitvi02g01019), the main transcription factor that regulates berry pigmentation, showed zero raw counts (TMM) in N. Steck at veraison, inducing a very high fold change value (< -8). On the other hand, at maturation, high and late expression of this gene (more than 700 TMM) was observed, with the gene becoming more highly expressed than in N. Branca and even matching the count values in N. Rosada.

Groups of DEGs from specific contrasts were isolated to target and identify meaningful differences in expression that could point to specific molecular components controlling the expression of the N. Steck berry phenotype (Table 1). The first group (1) corresponded to the intersection between (a) the genes that were negatively regulated during the transition from veraison to full maturation in N. Rosada and N. Branca but not in N. Steck and (b) the genes that were positively regulated in N. Steck compared with the other varieties within the maturation contrasts. The second group (2) represented the opposite case: the intersection between (a) the genes that exhibited constant expression or overexpression in N. Rosada and N. Branca but that were negatively regulated in N. Steck during the transitions of the maturation process and (b) the genes that were negatively regulated in N. Steck in comparison to the other varieties upon full berry ripeness (maturation stage).

**Table 1.** Enriched GO terms and KEGG pathways of the differentially expressed genes (DEGs) in the intersection groups. Group 1: a) Genes showing increased expression across stages in N. Steck and decreased expression in the other genotypes. b) Genes with constant expression in N. Steck across stages and decreased expression in the other genotypes. ab) Intersection of genes from a) and b). Group 2: a) Genes exhibiting increased expression across stages in N. Rosada and N. Branca and reduced expression in N. Steck. b) Genes with constant expression in N. Rosada and N. Branca across stages and reduced expression in N. Steck. ab) Intersection of genes from a) and b).

| 1 | DEGs | Gene Ontology - Biological Processes | KEGG Pathways |
| --- | --- | --- | --- |
| a | 282 | • response to auxin (GO:0009733);<br>• circadian rhythm (GO:0007623);<br>• endoplasmic reticulum to cytosol auxin (GO:0080162);<br>• auxin-activated signaling pathway (GO:0009734); | • vvi04075:<br>Plant hormone signal transduction |
| b | 607 | • response to biotic stimulus (GO:0009607);<br>• phenylpropanoid biosynthetic process (GO:0009699);<br>• dolichol-linked oligosaccharide biosynthetic process (GO:0006488) | • vvi00940:<br>Phenylpropanoid biosynthesis<br>• vvi00945:<br>Stilbenoid, diarylheptanoid and gingerol biosynthesis<br>• vvi00360:<br>Phenylalanine metabolism |
| ab | 100 | • transmembrane transport (GO:0055085) | • vvi04075:<br>Plant hormone signal transduction |
| 2 | DEGs | Gene Ontology - Biological Processes | KEGG Pathways |
| a | 1,343 | • translation (GO:0006412);<br>• plant-type secondary cell wall biogenesis (GO:0009834);<br>• regulation of DNA-templated transcription (GO:0006355);<br>• protoporphyrinogen IX biosynthetic process (GO:0006782);<br>• chlorophyll biosynthetic process (GO:0015995) | • vvi00940:<br>Phenylpropanoid biosynthesis;<br>• vvi00941:<br>Flavonoid biosynthesis;<br>• vvi00195:<br>Photosynthesis |
| b | 97 | • sterol biosynthetic process (GO:0016126);<br>• recognition of pollen (GO:0048544) | • vvi02010:<br>ABC transporters;<br>• vvi00073:<br>Cutin, suberin and wax biosynthesis;<br>• vvi00073:<br>Flavonoid biosynthesis |
| ab | 22 | • sterol biosynthetic process (GO:0016126);<br>• wax biosynthetic process (GO:0010025) | • vvi00073:<br>Cutin, suberin and wax biosynthesis; |

The first group represented the intersection between the 282 genes that had constant expression or extremely high expression in N. Steck and were negatively regulated in both N. Rosada and N. Branca during the berry development process (across stages) and the 607 genes more highly expressed in N. Steck than in both other varieties, consisting of 100 total genes. Genes of this intersection were enriched in the biological processes transmembrane transport (GO:0055085) and plant hormone signal transduction pathway (vvi0475), specifically genes encoding SAUR-related (Vitvi03g01365, Vitvi03g01367, and Vitvi08g00846) and ERF1/ERF2-related (Vitvi05g00715) enzymes, indicating disruption of auxin and ethylene processes in N. Steck berries.

The second group represented the intersection of the 1,343 genes that were negatively regulated in N. Steck and showed constant or extremely high expression in both N. Rosada and N. Branca during the berry development process (across stages) and another 97 genes less highly expressed in N. Steck than in the other varieties, consisting of 22 genes. This group included fewer genes than the first, although its enrichment provided interesting insights regarding negatively regulated processes in N. Steck. The enrichment of sterol and wax biosynthetic processes and cutin, suberin, and wax biosynthesis pathways indicated disruption of wax and cutin production and deposition in N. Steck berries. More specifically, the genes responsible for enrichment of the pathways are different isoforms of CER1, a gene responsible for cutin biosynthesis. The variance in expression patterns among ten specific genes was effectively validated through the RT-qPCR methodology, the results are visually depicted in Supplementary Figure S1.

#### Gene Ontology And Pathway Enrichment

All 6,247 DEGs were successfully annotated, and 2,732 different GO terms were retrieved. At veraison (Fig. 4A), DEG GO enrichment profiles were very dissimilar between contrasts, and N. Rosada showed more highly expressed genes with annotations for response to abscisic acid (GO:0009737) and lower expression of proteolysis genes (GO:0006508) than the other varieties. N. Branca showed an intermediate profile, with some terms enriched in the comparison with N. Rosada and N. Steck. Most of the enriched terms in N. Steck were related to upregulated genes, especially those associated with RNA polymerase II (GO:0006357, GO:0000978) and regulation of DNA transcription (GO:006355, GO:0001228). At the ripe stage (Fig. 4B), 4 enriched terms were upregulated in both N. Steck comparisons: L-phenylalanine ammonia-lyase activity (GO:0045448), polyketide biosynthetic process (GO:0030639), extracellular region (GO:0009607) and trihydroxystilbene synthase activity (GO:0016126). All the terms with downregulated enrichment between N. Rosada and N. Branca (NRM vs NBM) were also found to be downregulated between N. Steck and N. Branca (NSM vs NBM). The sterol biosynthetic process (GO:0016126) was equally downregulated in enrichment in N. Steck compared with the other genotypes.

**Figure 4.**
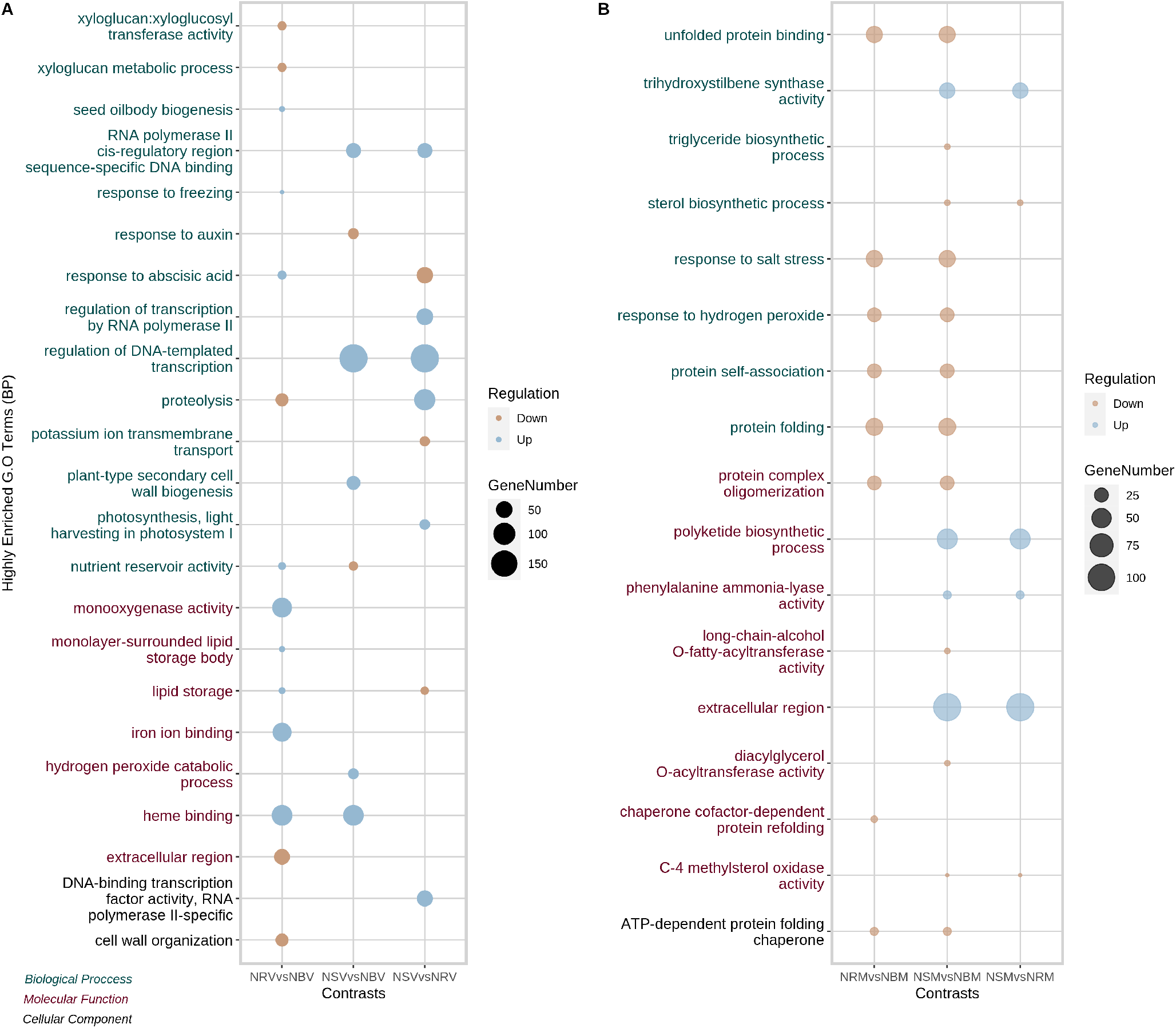
Scattergrams of enriched GO terms in expression contrasts between genotypes at both berry development stages. A) Veraison. B) Maturation.

Pathway enrichment analysis revealed fifteen different enriched KEGG pathways, seven at veraison (Fig. 5A) and twelve at maturation (Fig. 5B); most (11 of 13) of the reported pathways were enriched only in the N. Steck genotype. At veraison, in addition to the typical classes related to phenolic compound biosynthesis (phenylalanine, phenylalanine, flavonoid, flavonol, and flavone), the photosynthesis pathway (vvi00195) was also highly upregulated and galactose metabolism (vvi00052) downregulated in N. Steck compared with N. Rosada. Overall, plant hormone signal transduction (vvi04075) and phenylpropanoid (vvi00940) were the main pathways concurrently associated with the 1,544 genes upregulated in both N. Steck contrasts at veraison (Fig. 5A). For maturation comparisons, the most enriched pathway was protein processing in the endoplasmic reticulum (vvi04141), which was upregulated in N. Branca compared with the other varieties. The phenylpropanoid biosynthesis pathway remained upregulated in the N. Steck contrasts, but some of its branches, such as flavonoids (vvi00941) and flavones (vvi00944), became downregulated. The N. Rosada vs N. Branca DEGs were, in both stages, annotated as plant hormone signal transduction, phenylalanine metabolism and flavonoid biosynthesis genes.

**Figure 5.**
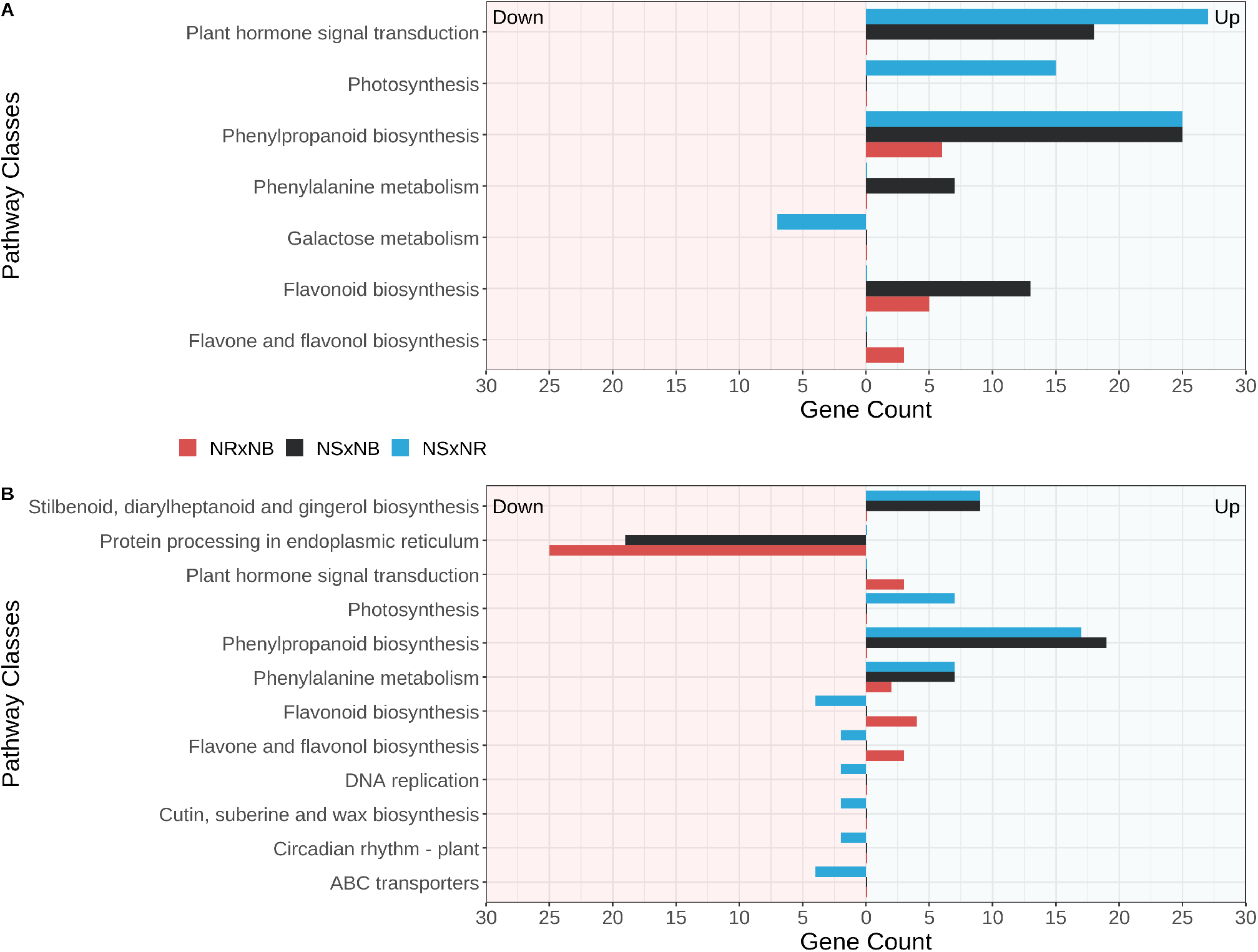
Enriched KEGG pathways between contrasts based on differentially expressed genes and their relative regulation. A) Veraison. B) Maturation.

### Coexpression Networks

The filtered expression data set was used to model a global coexpression network with the WGCNA approach. Pearson correlation coefficients were calculated and used as a similarity measure to model pairwise gene interactions. To fit the network to a scale-free topology, a *β* power of 14 was selected (scale-free topology model fit with R2*>*0.9 and mean connectivity of ≈ 540), and a corresponding dissimilarity matrix was calculated. The resulting network presented 39 groups of coexpressed genes with sizes ranging from 50 to 6,392 genes. To evaluate the importance of specific groups in the maintenance of gene interactions, we represented the WGCNA groups as nodes in a network (Fig. 6) and evaluated the degree and betweenness measures of nodes and edges, respectively (Supplementary Table S2). This representation of the coexpression network presented two central hub groups (1 and 2) with very discrepant degree values (20 and 22, respectively) compared with those of the others (mean degree of 6.2 of the network), as evidenced by the sizes of their nodes in Fig. 6. Additionally, 4 groups (17, 23, 24, and 33) did not show any connectivity with any other modules in the network.

**Figure 6.**
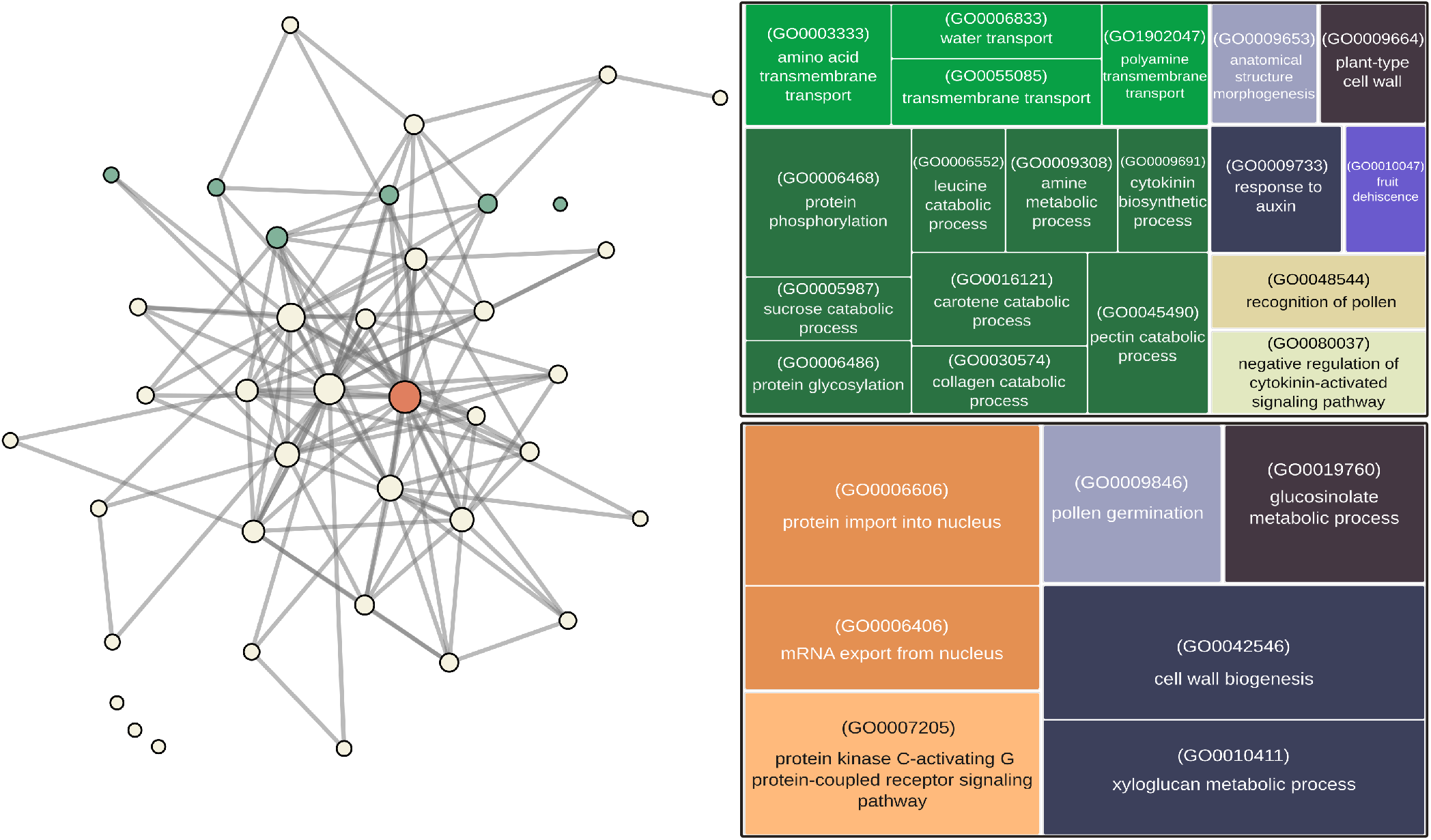
WGCNA global coexpression network and GO enrichment of nodes overrepresented by DEGs from the 1ab intersection (green) and 2ab intersection (orange). Node size represents the number of genes in the group, and edge width represents betweenness.

Our focus on the global coexpression network was identifying the modules composed of coexpressed genes that could be related to biological functions determining the phenotypic differences observed in the berries of the N. Steck genotype during its maturation process and, consequently, with its intensified russet characteristics. Therefore, modules that were enriched with the same intersection of DEGs described in Table 1 were sought within the network. Modules 4, 10, 13, 14, and 39 (green colored, Fig. 6 A) showed high representation (p value of Fisher’s test *<*0.05) of the DEGs from the **1ab** intersection; they were all closely positioned on the upper side of the network’s topology, and apart from module 39 (with 2 paths of distance), they all shared a direct edge with at least one other **1ab** enriched module. They also showed great proximity and direct connections with the two most central modules of the network (1 and 2). The GO enrichment of these **1ab** gene-enriched modules provided some insights into the biological processes and pathways driving the expression dynamics related to the intensified russet characteristics of the N. Steck berries, such as transmembrane transport (GO:0003333, GO:0006833, GO:0055085, GO:1902047), cell wall organization (GO:0009664), hormonal feedback related to auxin (GO:0009733) and cytokinin (GO:0009691, GO:0080037), fruit dehiscence (GO0010047) and many terms related to catabolism of different metabolites (GO:0006468, GO:0005987, GO:0006486, GO:0006552, GO:0009308, GO:0016121, GO:0030574, GO:0045490) and steroid biosynthesis (vvi00100) and the glycine, serine and threonine metabolism (vvi00260) pathways (Fig. 6 B).

The genes of the **2ab** intersection group had high representation in module 1 of the global network (orange colored), indicating a close relationship with genes that comprise the core of the network topology and therefore with the key genes controlling the physiology of berry development and maintenance. This was highlighted by module enrichment with general and basic biological processes such as the nucleocytoplasmic transport (vvi03013) KEGG pathway, which, in turn, is related to the transport of molecules in and out of the nucleus. This was further evidenced by the enriched GO terms protein import into nucleus (GO:0006606) and mRNA export from nucleus (GO:0006406) among the module’s genes. Other enriched terms worth mentioning are cell wall biogenesis (GO:0042546) and xyloglucan metabolic process (GO:0010411), which must be linked to the differential composition of N. Steck epidermis cells with suberin (Fig. 6 C).

The analysis of genotype-specific networks using the HRR methodology helped, in a more specific manner, to identify additional genes that are involved in the molecular control of the observed characteristics in N. Steck (Supplementary Fig. S2 E-F). The networks comprising genes coexpressed with genes from the intersection group **1ab** revealed two transcription factors of the MYB family (Vitvi18g00406, Vitvi01g00956) and the gene VvEXPA14 (Vitvi13g00172) as hubs in the N. Steck network. VvEXPA14 is associated with cell wall loosening and extension^15^; thus, it is likely one of the key mediators of the suppression of genes involved in softening of the peel in other N. Steck genotypes. In relation to N. Branca and N. Rosada (Supplementary Fig. S2 A-D), the networks revealed hub genes related to functions previously discussed, such as proteolysis (Vitvi13g01114), ABC transporters (Vitvi13g00638), and sterol biosynthesis (Vitvi13g00345).

Regarding the HRR networks including genes coexpressed with genes from intersection **2ab**, MYB4B (Vitvi04g01486) was the top hub gene in N. Steck. MYB4B has been previously reported as a repressor of phenylpropanoid biosynthesis, specifically in the production of low-molecular weight phenolic compounds^16^. In N. Branca, the annotated hubs identified were associated with processes related to the plasma membrane (Vitvi12g00410, Vitvi05g00360). In N. Rosada, the Vitvi04g00735 gene was identified as a top hub, which has been reported to be involved in the softening of the peel during maturation^17^, highlighting the importance of its repression in this process.

## Discussion

This study focused on the expression differences and coexpression of genes between three contrasting varieties of Niagara grapevine during two opposing stages of berry development. Divergent features were observed, in addition to the berry phenotype, at both the anatomical and molecular levels. The N. Rosada and N. Branca genotypes were the most similar, especially at the maturation stage; on the other hand, the N. Steck genotype showed the greatest differences, not only in berry skin structure and composition but also in expression levels and enriched biological functions and pathways, which could be associated with several elements of berry development and structure, such as transcription regulation, membrane and cell-wall dynamics, lipid production, auxin and cytokinin response, fruit dehiscence, biosynthesis of phenylpropanoid products and catabolism of many different compounds.

The N. Steck genotype results from a spontaneous somatic mutation and exhibits russet-like characteristics that are much more extreme than those reported in the few accounts of russet grapes (mainly ‘Shine Muscat’)^18^. An almost completely absent cuticle accompanied by multilayered peridermal scar tissue formation leads to the production of golden/brownish, thick, rough and impermeable skin, with thickened cell walls composed of lipid materials and no lignin. Except for the absence of lignin^19^, its structure is consistent with the usual anatomical descriptions of russet fruits in the literature^20^, such as in pears, apples and tomatoes. A striking difference of the russet phenotype in the N. Steck genotype, compared to that described in the literature, is its stable yearly expression in every berry of all clones, regardless of location, adverse environmental conditions and agronomic practices, indicating a genomic effect rather than a response to exogenous conditions.

The veraison stage marks the initiation of many regulatory changes in berry physiology that culminate in its maturation, primarily by a decrease in auxin biosynthesis, paired with an increase in abscisic acid production and sugar accumulation. Cell division ceases and gives rise to cell elongation (berry swelling), cell wall remission (skin softening) and secondary metabolite production and accumulation (pigments, flavor and aroma). This stage displayed the greatest differences in gene expression between our investigated varieties, with the greatest differences observed in comparisons with the N. Steck genotype. The heatmap (Fig. 3) of the 20 most DEGs shows that many of N. Steck’s genes at veraison, become non-DEGs at the maturation stage, indicating late and disrupted gene expression regulation, which is corroborated by the enrichment of several biological processes related to DNA to RNA transcription. VvUFGT and VvMYBA1, the main regulators of anthocyanin production, are among these late-expressed genes, showing no expression at veraison and counts as high as in N. Rosada at maturation, even though no anthocyanin pigmentation is observed in N. Steck berries. The phenylpropanoid pathway, responsible for the production of many secondary metabolites that enhance grape berry quality, was one of the most enriched pathways, with upregulated genes, in the N. Steck genotype. Nevertheless, the phenotype of N. Steck indicates that the phenolic products of this pathway, such as flavone and flavonoids, are also enriched, and anthocyanins are unable to accumulate in the berry tissue.

Network groups enriched with the **1ab** (Table 1) intersecting genes allowed the identification of many enriched GO terms related to the catabolism of different types of metabolites, such as sucrose, carotene, collagen, pectin, amine and leucine, indicating that these upregulated genes are correlated with genes encoding proteins associated with molecular catabolism. Furthermore, these groups were also enriched in the fruit dehiscence term, pointing to a direct correlation between these DEGs and the expression of other genes regulating berry development. At the maturation stage, GO and pathway enrichment showed many terms equally enriched in N. Steck and N. Rosada when they were compared to N. Branca, revealing more similar expression profiles between them at this stage, which is expected since N. Steck is a somatic mutation of N. Rosada, although they have very distinct berry phenotypes and N. Steck still shows more DEGs. This result supports the hypothesis of late ripening and disrupted circadian rhythm in N. Steck, which was also a downregulated pathway in this genotype at this developmental stage.

Complex phytohormone interplay and cross-talk mediate each stage of berry development, and even slight imbalances in these processes can interfere with the achievement of a complete ripening state^21^. Our data showed great upregulation of the plant hormonal signal transduction pathway in N. Steck at veraison, especially compared with that in N. Rosada. The enrichment of this pathway resulted from the differential expression of SAUR and ERF enzyme-encoding genes, which are related to auxin and ethylene signalization and biosynthesis, respectively. Auxin has an important role in determining early berry structure by promoting cell division; however, the remission of its content and participation at veraison are necessary for the initiation of berry ripening, and extended periods of auxin presence greatly delay berry ripening^22^. The higher expression of SAUR-related genes indicates that auxin-related processes are still occurring in N. Steck, but not in the other varieties. These processes should be related to the formation of the epicarp layer observed on the skin in N. Steck. Considering that this layer is present on the entire surface of the berry skin, its development must require a high and constant rate of cell division mediated by auxin and, consequently, disruption of the expected development period, although our data are not sufficient to determine whether auxin levels are truly different in N. Steck and if this process or another antecedent molecular process is disrupted. Functional enrichment of network groups enriched with the **1ab** intersection genes also supported a relationship between these DEGs and the expression of genes annotated for response to auxin. With regard to ethylene, the reported DEGs of ERF’ also indicate a disturbance in its signaling and biosynthesis processes. Even though grape berries are not climacteric, there are reported ethylene peaks during the onset of veraison^23^, associations with anthocyanin production^24^, and therefore, a secondary role of this phytohormone in the berry development of N. Steck; perhaps this effect is related to the intense increases in VvUFGT and VvMYBA1 expression at maturation. The **1ab** enriched groups in the networks were also associated with two terms related to the cytokinin process. Investigations on the participation of this phytohormone in berry development demonstrated that exogenous applications or extended periods of constant presence of cytokinin in the berry tissue can lead to a delay in maturation, along with a reduction in anthocyanin accumulation and an increase in cuticle content^25,26^. These results demonstrated that the N. Steck phenotype is not linked to a specific hormone imbalance but to major interference in the whole hormonal interplay that mediates the ripening process.

The most evident variances in N. Steck berries were those related to plasma membrane and cell wall structure, composition and transport. These variances were observed in every performed analysis, which gave us insights into different aspects of their emergence. The observed epicarp formation was accompanied by the thickening of cell walls in the N. Steck genotype and irregular, almost absent, formation of the cuticle layer. Yang et al. (2023)^27^ demonstrated that the removal of epicuticular wax from grape berry skin accelerated berry weight loss, chlorophyll degradation, browning, softening and polyphenol biosynthesis, suggesting a protagonist role of the gene HMGR2 (Vitvi18g00043) in wax reconstitution after skin removal. Apart from berry softening, these consequences are very similar to the N. Steck berry characteristics, and interestingly, the HMGR2 gene was more highly expressed in N. Steck at the maturation stage than in the other varieties, indicating that dysfunction of wax biosynthesis must be causing the absence of the cuticle layer, possibly triggering a reconstitution process. It also seems that the regeneration process of the cuticle layer is not successful, which could lead to an increase in water loss of the cells, triggering a response involving the expression of genes (mainly of the MYB family) involved in the biosynthesis and deposition of suberin and other molecules in the skin. The network groups enriched with **1ab** intersection genes allowed the identification of specific molecules not present in the intersection annotation (Table 1) that were transported across the plasmatic membrane, such as amino acids, polyamines and water. This accumulation, mainly of lipids, prevents the cells from bursting and dying and causes the brownish, rough and thick aspects of N. Steck skin, causing it to resemble root/bark-like tissue. This hypothesis is also supported by the enrichment of sterol and wax biosynthetic terms and pathways among the **2ab** intersection genes, i.e., those that are downregulated in N. Steck compared with the other varieties. This intersection group was composed, among others, of genes encoding CER family proteins, which are associated with the production of epicuticular wax, pointing to a deficiency in the biosynthesis of fatty acids in N. Steck, which play important roles in berry protection against biotic agents, such as insects and microorganisms, and in tolerance to abiotic stresses, such as water permeation resistance, salt tolerance, dust prevention and sunlight blocking. Additionally, cuticular wax enhances berry attractiveness for consumption because it promotes glossiness, brightness and luster and is closely related to fruit development and postharvest storage quality. In russet-susceptible apple varieties, increased cell divisions during fruit development increase epidermal strain, causing microcracks in the cuticle layer, which leads to its replacement with suberin and, upon contact with the atmosphere, brownish patches on the epidermal layer^28^. This mechanism could explain the emergence of the N. Steck berry phenotype, but instead of occurring only in the cuticle layer, which is almost completely absent, these microcracks also occur in the developed epicarp layer, as shown by scanning electron microscopy, and must cause a signal that induces intense thickening of the skin cell walls with lipids, mainly suberin, and, consequently, impermeabilization of the berry. Last, network module 2 also showed the presence of multiple orthologs of both the CER1 (Vitvi18g00529, Vitvi06g01456) and MYB41 (Vitvi10g00329, Vitvi19g00306, Vitvi02g00725, Vitvi14g01987, Vitvi16g00098, Vitvi11g01323) genes and could represent the interface point of cuticle remission, suberin deposition and the usual expression changes that control berry development and maturation.

## Methods

### Plant Material

‘Niagara Branca’, ‘Niagara Rosada’, and ‘Niagara Steck’ (Fig. 7a) grapevines were cultivated in the same vineyard and with the same viticulture management practices at the experimental fruit station of the Agronomic Institute, Jundiaí, Brazil (latitude 23°06” S, longitude 46°56” W). The vines were sustained in an espalier system and pruned in August every year, leaving one or two buds per branch. The plants were grafted onto the “IAC 766 Campinas”rootstock in 2008 with a spacing of 2.0 m x 1.0 m between rows and plants, and each accession was represented by three clonally propagated plants. All plants of the collection were routinely screened for diseases and pathogen infection.

**Figure 7.**
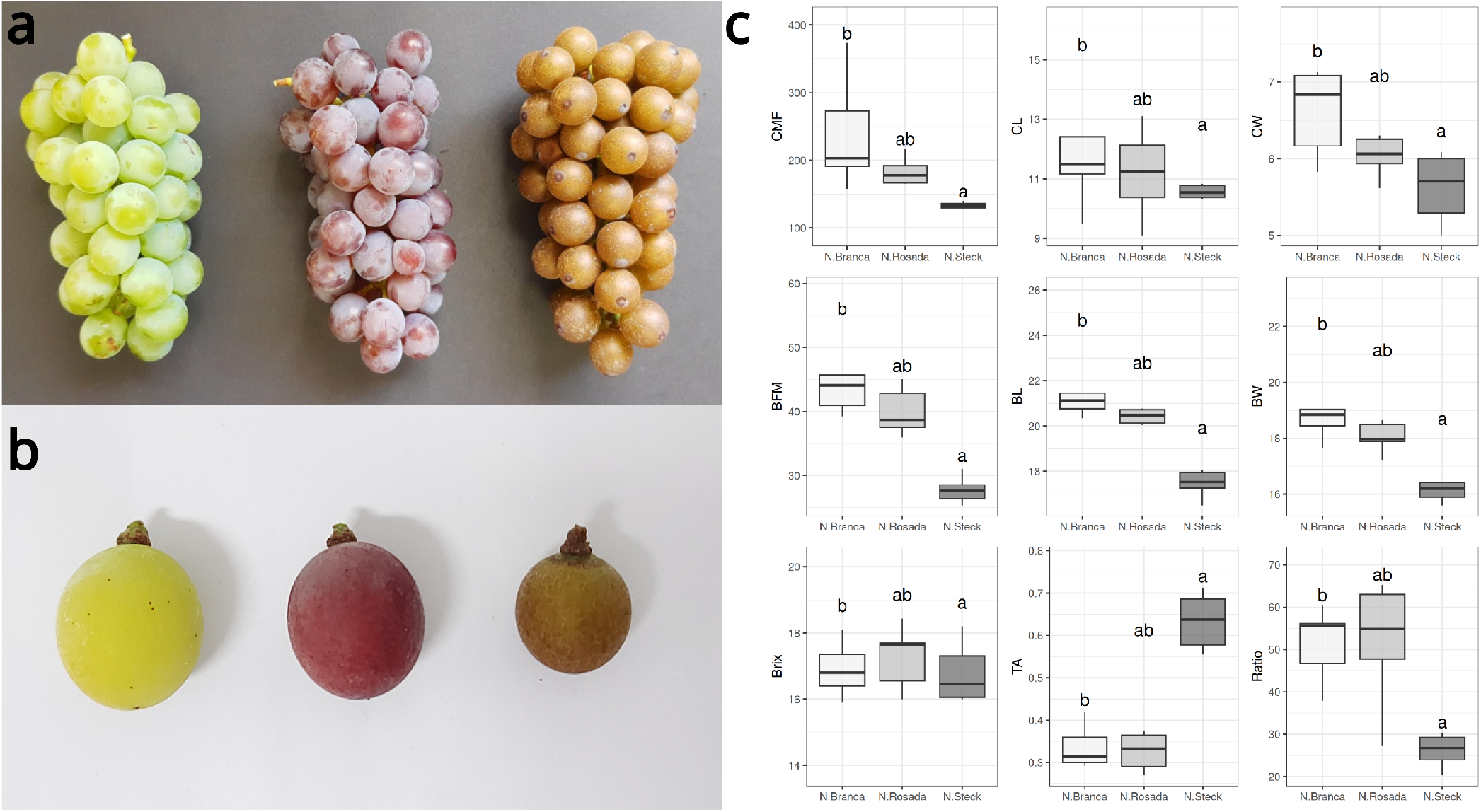
a) Ripe clusters from N. Branca, N. Rosada and N. Steck. b) Berry size and pigmentation comparison. c) Tukey’s test of phenotypic measurements from 2013 to 2021. CFM: Chunk fresh mass, CL: Chunk length, CW: Chunk width, BFM: Berry fresh mass, BL: Berry length, BW: Berry width, Brix: Brix degree, TA: Titratable acidity, Ratio: Maturation Ratio

Fruit samples were harvested at approximately 10 a.m. in 2020 at the veraison (65 days after full bloom - DAFB) and ripe (90 DAFB) stages throughout the growing season. On both sampling dates, three independent biological replicates (plant individuals) of each genotype, positioned in the same row within the vineyard, were selected, and from each replicate, 3 clusters were chosen. Ten berries from each cluster were picked, from both row sides, and those collected at the veraison stage were selected to form a pool of green-to-slightly pigmented berries (N. Rosada and N. Steck) in order to obtain the most reliable representation of the phenotype at this stage. Berries were immediately frozen with dry ice and stored at −80 °C.

### Berry Structural Analysis

#### Light Optical Microscopy

For the anatomical analysis of the skin region, the berries at both sampled developmental stages were fixed in FAA (formaldehyde, acetic acid, ethanol 50%/ 1:1:18 v/v) for 24 hours^29^ and stored in 70% ethanol. The material was dehydrated in an ethanol series and embedded in hydroxyethyl–methacrylate (Historesin® Leica) following the manufacture’s recommendations. Cross-sections 0.5 *μ*m thick were made with the aid of a Microm HM 340E rotary microtome (Thermo Fisher Scientific Inc., Waltham, Massachusetts, USA). The slides were stained with 0.05% toluidine blue in citrate buffer (pH = 4.5)^30^ and finally mounted in EntellanÂ® synthetic resin (Merck KGaA, Darmstadt, Germany). Images were captured with a digital camera (Olympus DP71) coupled to an Olympus BX51 optical microscope (Olympus Optical Co., Ltd., Japan).

#### Scanning Electron Microscopy

For micromorphological analysis of the epicarp surface, the berries were fixed in FAA (formaldehyde, acetic acid, 50% ethanol; 1:1:18 v/v) for 24 hours^29^ and stored in 70% ethanol. Subsequently, they were dehydrated in a graduated ethanol series, dried by the critical point method using CO_2_ in a CPD-030 apparatus, mounted and metallized with gold (exposure time 200 s on a Balzers SCD-050 sputter coater, ONLINK Technologies GmbH, Germany). The observations and images were obtained using a JEOL JSM 5800 LV scanning electron microscope (SEM) at 10 kV with a digital camera attached (GenTech Scientific Inc., New York, USA).

#### Histochemical Analysis

Berry samples were fixed, dehydrated, included and photographed as described above. To detect the main chemical compounds present in the berry skin of different grape varieties, we used the following histochemical tests: ruthenium red for acidic mucilage^29^, ferric chloride for phenolic compounds^29^, Sudan III for total lipids^31^ and Nile Red fluorochrome for lipids and cutin, under a DAPI-LP filter (ex 377/50 nm; em 447/60 nm)^32^.

### RNA Extraction, Library Construction and Sequencing

Frozen pulp and skin of each picked berry were ground into a powder with a pestle and liquid nitrogen, and the seeds were disposed of. RNA molecules were isolated and purified from the powdered tissues with the protocol described by Reid et al. (2006)^33^. RNA sample integrity was determined by 1% (w/v) agarose gel electrophoresis, and purity and concentration were determined by a NanoVue spectrophotometer (GE Healthcare, Chicago, IL, USA). An RNA pool for each biological replicate was generated by mixing the samples extracted from the picked berries, and 1 *μ*g of each pool was subsequently used to build distinct cDNA libraries with the TruSeq Stranded mRNA Library Prep (Illumina, San Diego, CA, USA) kit. Library concentrations were verified by qPCR (Illumina SY-930-1010 protocol), and library composition and purity were verified by an Agilent 2100 Bioanalyzer (Agilent Technologies, Palo Alto, CA, USA). Next-generation sequencing was performed on a NextSeq 500 instrument (Illumina, San Diego, CA, USA).

### Bioinformatics

#### Genome Mapping and Transcriptome Assembly

Low-quality reads were identified by FastQC^34^ and filtered on the basis of a minimum Phred score of 30 with Trimmomatic^35^. High-quality reads were mapped to the grapevine 12X.2 reference genome assembly along with its structural annotations Vcost.V3^36^ with STAR^36^, and reference and novel transcripts were assembled by StringTie^37^ and merged to create a transcriptome file. Transcriptome quality assessments were performed by benchmarking the assembled transcripts against the universal single-copy orthologs of the Viridiplantae clade with BUSCO^38^ with default parameters. Reads were mapped to the transcriptome with Bowtie 2^39^, and transcript abundances were estimated with Salmon^40^.

#### Differential Expression Analysis

Transcript counts were aggregated at the gene level with the package tximport^41^ for R. The EdgeR R package^42^ was used to normalize counts by the Trimmed Mean Method (TMM), filter weakly expressed genes by the *filterByExpr* function, with a minimum count set to 10, and perform differential expression analysis. Nine combinations between the samples were established as contrasts (Table 2) and were grouped according to genotype (different genotypes at the same berry development stage) or grape development stage (the same genotype at different stages). A logFC absolute value ≥ 2 and a false discovery rate (FDR) ≤ 0.05 were used for the identification of DEGs. We selected the top 20 genes with the largest fold change values in the comparisons between varieties and represented the fold change measures, in such comparisons, using a heatmap coupled with a dendrogram constructed based on a complete hierarchical clustering analysis performed on dissimilarities calculated with Pearson correlations between the fold change values.

**Table 2.** Expression contrast groups for the genotype and berry development stage comparisons.

| Contrast | Genotype (baseline) | Genotype | Berry Stage |
| --- | --- | --- | --- |
| NRV vs NBV | N. Rosada | N. Branca | Veraison |
| NSV vs NBV | N. Steck | N. Branca | Veraison |
| NSV vs NRV | N. Steck | N. Rosada | Veraison |
| NRM vs NBM | N. Rosada | N. Branca | Maturation |
| NSM vs NBM | N. Steck | N. Branca | Maturation |
| NSM vs NRM | N. Steck | N. Rosada | Maturation |
| NSM vs NSV | N. Steck | N. Steck | Veraison/Maturation |
| NBM vs NBV | N. Branca | N. Branca | Veraison/Maturation |
| NRM vs NRV | N. Rosada | N. Rosada | Veraison/Maturation |

#### Annotation, Gene Ontology and Pathway Enrichment

The functional annotation of transcripts was performed considering (i) *Vitis vinifera* V.3 gene annotations from the Ensemble Plants database^43^ and (ii) comparative alignments of StringTie-assembled sequences against UniprotKB/SwissProt using diamond software^44^. Only hits with e-values *<* 10^−5^ and percentages of identical matches > 80% were accepted. Gene Ontology (GO) enrichment analysis of DEGs and coexpression network modules was performed using the topGO R package^45^. Reported GO terms with adjusted p values *<* 0.005 were considered significantly enriched, and term redundancy was assessed with Revigo^46^. The ClusterProfiler^47^ package for R was used for the identification of enriched KEGG pathways between the contrasts, enlisting the same sets of DEGs and coexpressed genes as in the gene ontology analysis.

### Coexpression Network Analysis

A global gene coexpression network was built with the WGCNA R package^48^ using normalized transcription counts based on a *β* soft-threshold power defined according to the lowest mean connectivity and a minimum linear regression model fitting index (R^2^) of 0.9. Network modules, with a minimum of 30 genes, were determined by transforming adjacency matrices into topological overlap matrices (TOMs) and merged when coexpression similarities exceeded 80%. The network modules numbers are available in the supplement (Fig. S1). Furthermore, additional networks were built for each genotype using the highest reciprocal ranks (HRR) method^49^ based on the expression of genes belonging to nodes of the WGCNA global network enriched with genes from the analyzed intersection groups of DEGs.

### Quantitative Real-time Polymerase Chain Reaction (RT-qPCR)

Ten transcripts were randomly selected to confirm the RNA-Seq expression profiles by RT-qPCR. The selection was based on transcripts with high logFC values, low p values and high absolute expression counts. A Quantitec Reverse Transcription Kit (Qiagen, Hilden, Germany) was used to perform reverse transcription of the extracted RNA samples. Primers with amplicon lengths ranging from 80 to 200 kb, a 60°C annealing temperature and a 20 bp primer length were applied to the selected transcripts with the Primer3Plus platform^50^(Supplementary Table S4). Amplifications were performed in triplicate with SYBR Green Supermix (Bio-Rad, Hercules, CA, EUA) fluorescence on a Bio-Rad CFX384 Touch Real-Time Detection System. Detected expression levels were normalized using the comparative CT method (∆∆CT method), and *Vitis vinifera* ACTIN1 and AP47 were used as reference genes.

## Acknowledgments (not compulsory)

The authors gratefully acknowledge the Coordenação de Aperfeiçoamento do Pessoal de Nível Superior (CAPES) for the fellowships awarded to GN (513247/2020-00), GO (479802/2020-00) and DG (140174/2021-4); the Fundação de Amparo à Pesquisa do Estado de São Paulo (FAPESP) for Ph.D. fellowships awarded to AA (2019/03232-6) and financial support (FAPESP, 2020/12938-7); and the Conselho Nacional de Desenvolvimento Científico e Tecnológico for financial support (CNPq, 404041/2021-3).

## Author contributions statement

GN and GO conducted and analyzed the RNA-Seq experiments and wrote the manuscript; AA assisted in the result analysis and manuscript writing; DG and SCG performed the LOM and SEM analysis; MM maintained the germplasm collection and performed the phenotypic analyses; and AS conceived the project. All authors reviewed, read and approved the manuscript.

## Additional information

The authors declare no competing interests.

