## Supplementary Figures for "Uncovering the molecular signatures of russet skin formation in Niagara grapevine (*Vitis vinifera x Vitis labrusca*)"

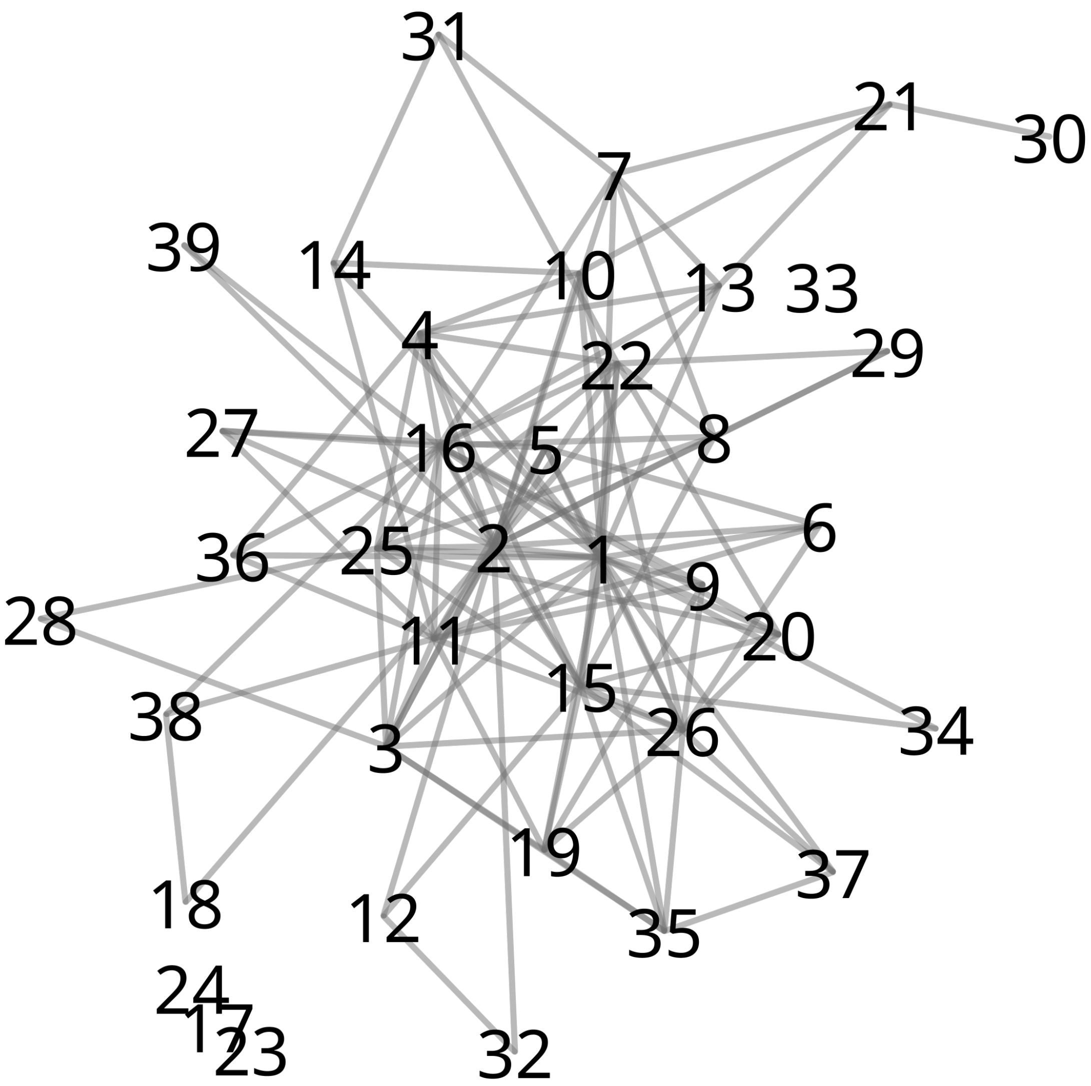


**Supplementary Figure S1:** WGCNA global co-expression network. Vizualization of the global expression network showcasing modules denoted by their respective module number.

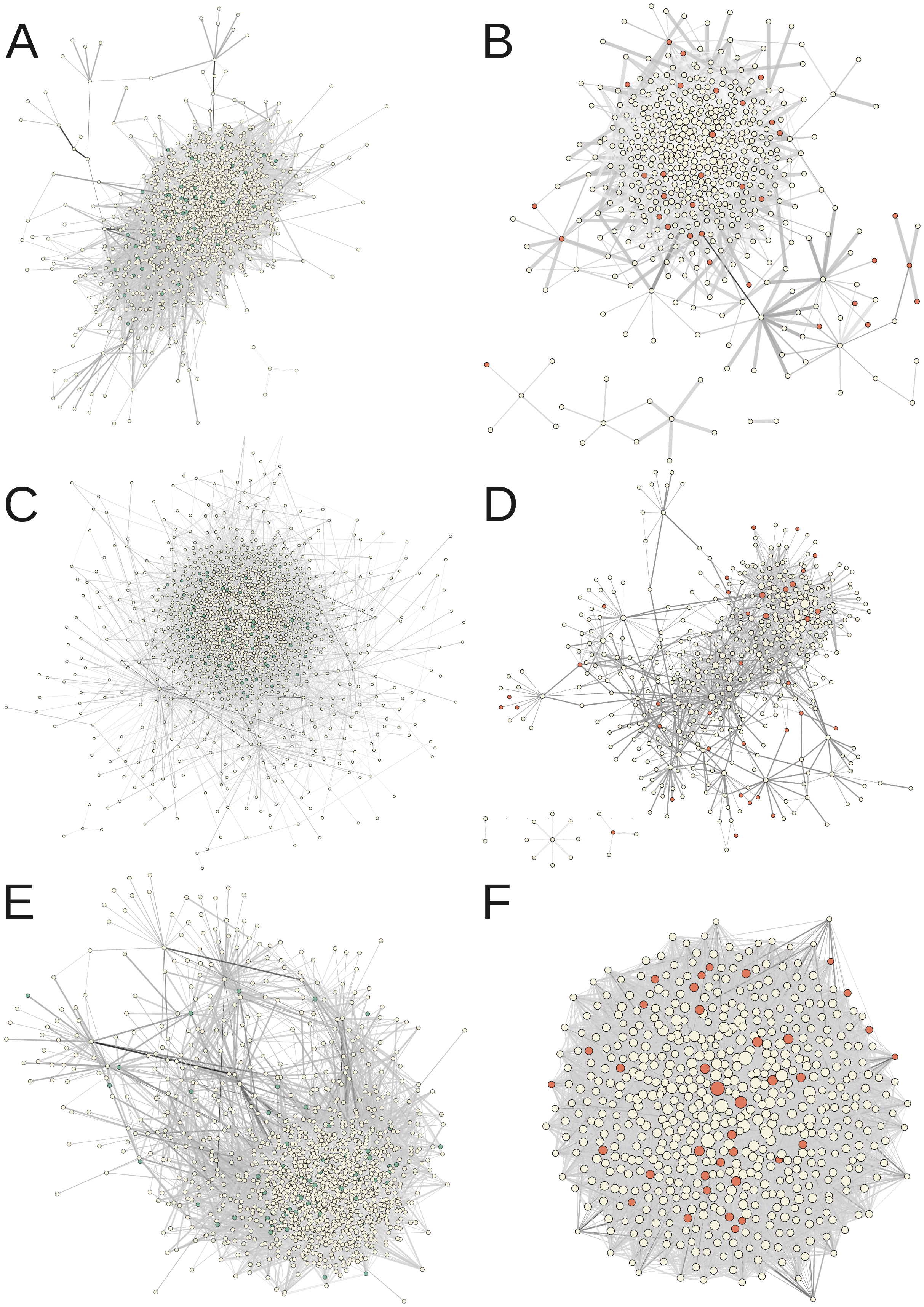


**Supplementary Figure S2:** Genotype-specific network analysis with the Highest Reciprocal Rank (HRR) method. A-B: Niagara Branca. C-D: Niagara Rosada. E-F: Niagara Steck. The network highlights green modules corresponding to genes from the **1ab** intersection group, and orange modules corresponding to genes from the **2ab** intersection group.

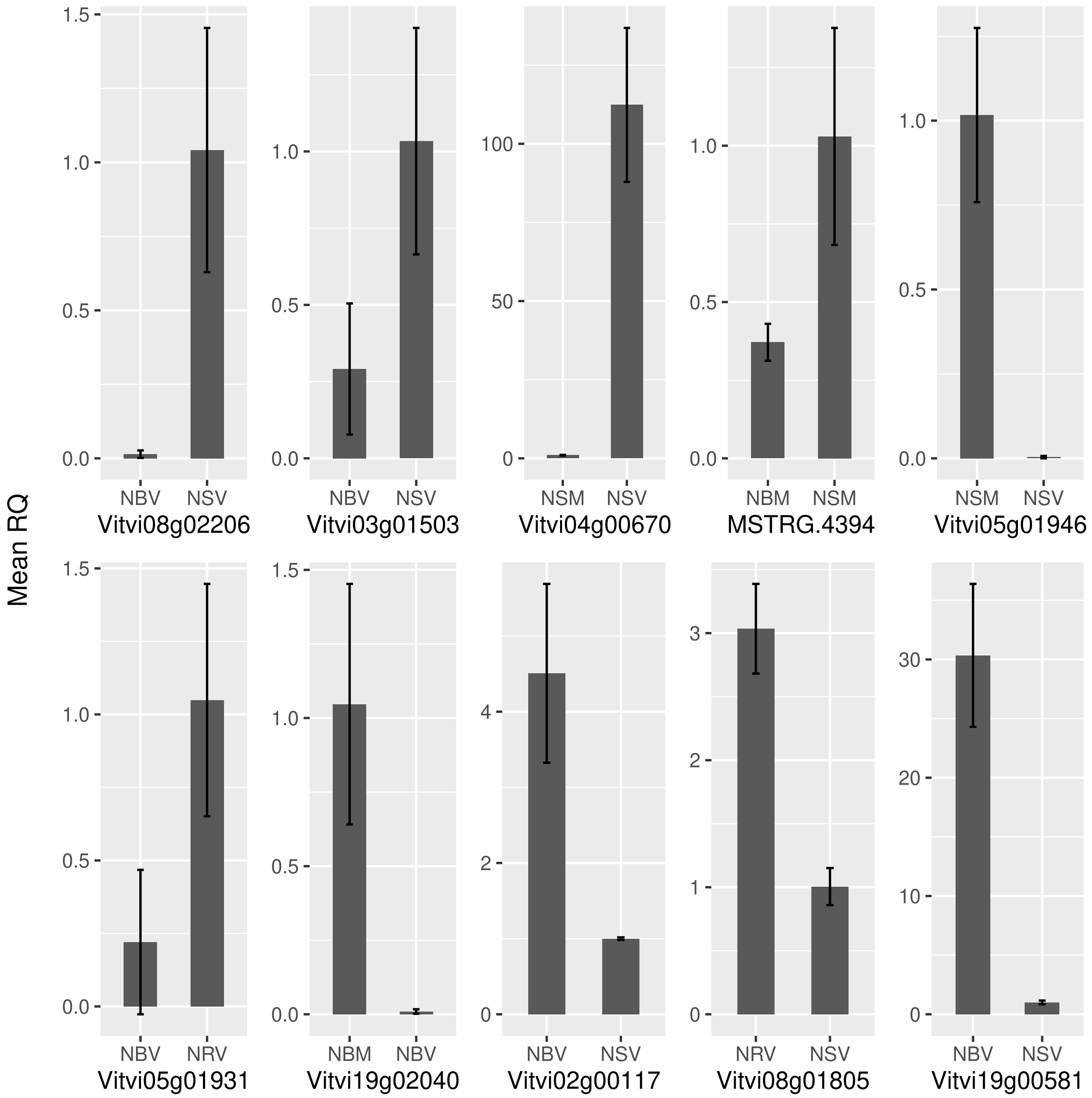


**Supplementary Figure S3:** Validation of the results from differential expression analysis. RT-qPCR technique was conducted using ten differentially expressed genes selected at random. Their corresponding cycle quantification (Cq) values were utilized to apply the ΔΔCq method, thereby determining the Relative Quantification (RQ) values. Statistical significance of the analysis was evaluated using the Student's t-test.
